# Tri-Modality Representation Learning for Molecular Property Prediction

**DOI:** 10.64898/2026.09.22.753673

**Authors:** Anyin Zhao, Jing Li

## Abstract

Accurate molecular property prediction requires effective molecular representations that can describe a molecule from multiple complementary perspectives. Existing deep learning approaches typically use SMILES strings, two-dimensional molecular graphs, or three-dimensional conformations as inputs. These representations capture different aspects of molecular information: SMILES encodes a sequential description, molecular graphs describe atom–bond connectivity, and 3D conformations provide spatial information from atomic coordinates. However, these modalities are often learned separately or merged with low effective fusion operations, which may not sufficiently capture interactions among sequential, topological, and geometric features. In this work, we propose Tri-Modality Cross-Attention (TMCA), a multimodal framework that integrates 1D SMILES, 2D molecular graphs, and 3D molecular conformations for molecular property prediction. TMCA uses pretrained encoders for the 1D and 3D branches, with a SMILES-based Transformer for the 1D branch and a recent conformation-aware pretrained model for the 3D branch. At the same time, a trainable 2D graph encoder is designed to support modality fusion. All the parameters from the pretrained models are frozen to make overall training more efficient. We evaluate TMCA on four datasets (*i.e*., BBBP, BACE, ClinTox, and HIV) from the MoleculeNet database for classification tasks and compare its performance with state of the art competing approaches. Experimental results show that TMCA achieves the best overall performance, demonstrating the value of multi-modality integration for drug property prediction and the effectiveness of our cross-attention based fusion strategy. The source code of TMCA is available at: https://github.com/Ay-Zhao/TMCA

## 1 Introduction

Traditional drug discovery is costly, time-consuming, and associated with high failure rates [14]. To reduce resource-intensive experimental screening, computational methods have been widely studied for molecular property prediction. In this paper, we focus on deep-learning-based molecular property prediction, which aims to help researchers prioritize molecules for further validation [18, 29, 6, 8].

Accurate molecular property prediction depends on effective molecular representations. Molecules can be described from multiple views, including 1D SMILES strings, 2D molecular graphs, and 3D conformations. SMILES strings represent molecules as symbolic sequences [20]. Molecular graphs encode atom– bond connectivity. And 3D conformations provide geometric information through atom types and atomic coordinates. Accordingly, Transformer-based models have been used for SMILES representation learning [15, 2, 17]. Graph neural networks have been widely adopted for molecular graphs [4, 25]. And recent 3D models such as Uni-Mol learn conformation-aware representations from molecular geometry [28].

These modalities provide complementary information. SMILES-based models are effective at capturing global sequential patterns. Graph-based models learn local topological structures. And 3D conformation-based models encode spatial relationships that are not explicit in 1D or 2D representations. However, using these modalities independently, or combining them with simple fusion operations such as concatenation, may not fully exploit complementary information provided by all three types of features.

TMCA extends our previous work named DMCA [27], which was based on integration of two modalities (*i.e*., molecular graph and SMILES). TMCA strategically integrates sequential, topological, and geometric modalities by using a trainable 2D graph branch as the central query modality. Specifically, TMCA uses pretrained encoders for the 1D SMILES and 3D conformation branches, and train a 2D graph encoder from scratch. Two graph-guided cross-attention modules are then used to retrieve complementary sequence and geometry information. The fused representations are then combined for downstream drug property prediction. We evaluate TMCA on four datasets (*i.e*., BBBP, BACE, ClinTox, and HIV) from the MoleculeNet database [22] for classification tasks and compare its performance with state of the art competing approaches. Results show that TMCA achieves best overall performance, demonstrating the value of multi-modality integration for drug property prediction and the effectiveness of our cross-attention based fusion strategy.

The rest of the paper is organized as follows. We introduce some related work in Section 2. The details of the proposed approach are discussed in Section 3. After discussing the experimental setup in Section 4, comparison results including results from ablation studies are presented in section 5. We conclude our paper with some discussions at the end.

## 2 Related Work

Recent studies on molecular property prediction mainly differ in how molecules are represented and how different molecular views are learned. In this section, we briefly review approaches relying on 1D SMILES strings, 2D molecular graphs and 3D molecular conformations individually, as well as some approaches that are based on multimodal molecular representation learning.

### 2.1 1D SMILES-based Molecular Representation

SMILES strings represent molecules as linear sequences [20], making them suitable for sequence models originally developed in natural language processing. Transformer-based models [15] have been widely adopted for SMILES representation learning because self-attention can capture long-range dependencies among tokens. SMILES-BERT [17] and ChemBERTa [2] used masked language modeling for molecular pretraining, while ST [7] adopted a Transformer encoder– decoder framework for learning from unlabeled SMILES strings. Other methods, such as X-Mol [24] and Mol-BERT [11], further explored self-supervised objectives for molecular sequence learning. Although SMILES-based models can benefit from large-scale pretraining and global sequence modeling, they do not directly encode molecular topology or 3D geometry. Furthermore, unlike natural languages, the token sequences in the SMILES format lack semantic meaning, which hinder the effectiveness of Transformer-based language models.

### 2.2 2D Graph-based Molecular Representation

Molecular graphs provide a natural representation of chemical structures, where atoms are modeled as nodes and chemical bonds are modeled as edges. Graph neural networks (GNNs) have therefore been widely used for molecular property prediction by propagating information along atom–bond connections [4, 25]. To improve graph representation learning, several studies have introduced self-supervised or contrastive pretraining strategies. Hu et al. [8] proposed node-level and graph-level pretraining tasks to improve molecular GNNs. MolCLR [18] applied contrastive learning to molecular graphs using graph augmentations. These methods are effective at capturing local topology and substructure patterns, but graph message passing can still have difficulty modeling long-range molecular interactions, especially when deeper GNNs suffer from over-smoothing [12].

### 2.3 3D Geometry-aware Molecular Representation

Beyond 1D sequences and 2D graphs, 3D molecular conformations provide geometric information through atom coordinates. This information can be important for properties related to molecular shape, spatial arrangement, and interatomic distances. Several recent methods incorporate 3D information into molecular representation learning. GEM [3] introduces geometry-based pretraining to capture spatial molecular knowledge. Uni-Mol [28] further develops a Transformer-based framework for 3D molecular representation learning from atom types and coordinates. These methods demonstrate the usefulness of geometric information, but using 3D features alone may overlook complementary sequential and topological signals.

### 2.4 Multimodal Molecular Representation Learning

Multimodal learning aims to combine complementary information from different data views [1, 23]. For molecular property prediction, many multimodal studies have focused on combining SMILES and 2D graph representations. For example DVMP [29] learned coordinated representations from SMILES and molecular graphs, while GraSeq [5] fused graph and sequence features through a gating mechanism. Our previous work, DMCA [27], further explored graph–SMILES fusion using a dual-modality cross-attention mechanism. MSSGAT [26] combines molecular graphs, substructure information, and fingerprint-related features. More recent work has started to incorporate 3D geometry together with other molecular views. For instance, GraphMVP [13] used both 2D topology and 3D geometry for self-supervised molecular pretraining.

TMCA is a natural extension of our previous work DMCA [27], by adding the third modality using 3D geometry information on top of the 1D and 2D modalities. Although some most recent approaches (*e.g*., SGGRL [19]) have explored this direction, TMCA is different in how it performs the integration. It uses a trainable 2D graph branch as the central query modality and retrieves complementary information from frozen pretrained 1D sequence and 3D geometric encoding through graph-guided cross-attention. This design provides a simple and extensible way to integrate pretrained and non-pretrained molecular representations.

## 3 Methods

In this section, we present Tri-Modality Cross-Attention (TMCA), a hierarchical neural architecture designed for joint molecular representation learning across three complementary modalities. TMCA extends our previous dual-modality cross-attention framework [27] by incorporating 3D molecular conformations. The central idea of TMCA is to improve molecular property prediction by integrating sequential, topological, and geometric information through Transformer-based multimodal fusion. Specifically, TMCA combines representations from 1D SMILES strings, 2D molecular graphs, and 3D molecular conformations using graph-guided cross-attention modules to construct a unified molecular embedding.

The overall architecture of TMCA (Fig. 1(a)) consists of four major components: (i) a Transformer-based SMILES encoder that takes 1D sequential SMILES strings as its input and captures global sequence-level molecular patterns; (ii) a GNN-based graph encoder that processes 2D molecular graphs and learns local atom–bond connectivity information; (iii) a Transformer-based 3D conformation encoder that produces conformation-aware molecular representations based on atom types and atomic coordinates, to capture geometric information; and (iv) two cross-attention fusion encoders that integrate modality-specific embeddings and generate the final molecular representation for downstream applications.

**Fig. 1.**
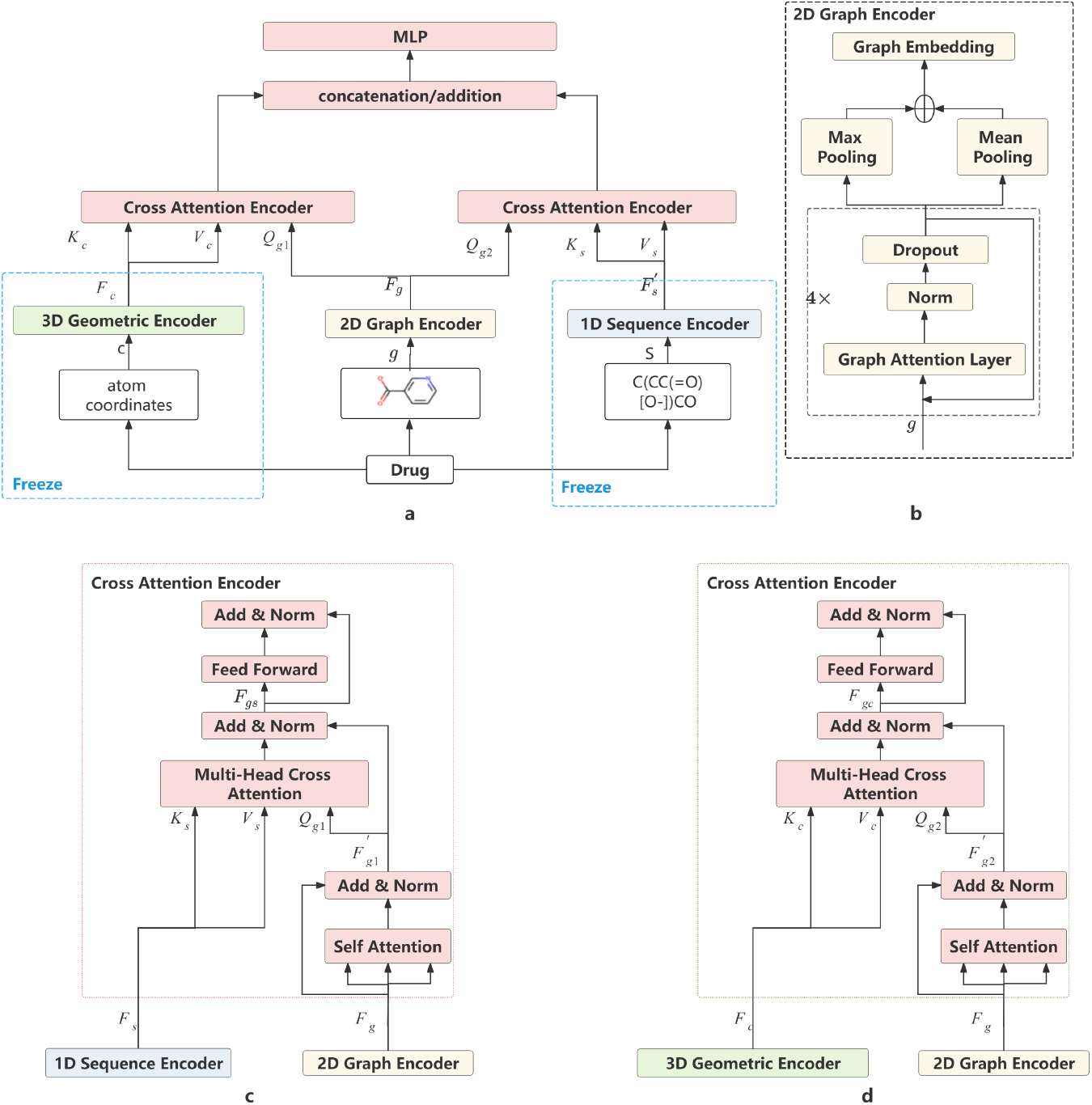
Workflow of the proposed TMCA model. (a) Overall tri-modality architecture using 1D SMILES, 2D graphs, and 3D conformations. (b) The 2D graph encoder. (c) The graph-guided sequential cross-attention encoder. (d) The graph-guided geometric cross-attention encoder

A key design of TMCA is to use the 2D graph representation as the central query modality in the cross-attention modules. The 1D sequence encoder and 3D geometric encoder are pretrained and provide stable modality-specific representations. The trainable 2D graph encoder learns to adaptively retrieve useful information from both pretrained branches. This design enables effective fusion between pretrained and non-pretrained encoders. It also reduces the computational cost compared with fully connected pairwise multimodal attention because TMCA only performs graph-guided attention from 2D to 1D and from 2D to 3D. After cross-attention, the fused embeddings are combined through a simple late-stage fusion operation, such as concatenation or addition, and then passed to an MLP predictor. This modular design makes TMCA easy to extend to additional molecular modalities by introducing new graph-guided cross-attention branches.

### 3.1 1D Sequence Encoder

SMILES provides a compact linear notation for describing molecular structures [20]. In the 1D branch of TMCA, each SMILES string is treated as a token sequence and encoded using a Transformer-based molecular language model. Before being passed into the encoder, the SMILES string is tokenized using the Byte-Pair Encoding (BPE) tokenizer adopted in ChemBERTa [2]. This tokenizer, implemented through the HuggingFace tokenizers library [21], combines character-level and substructure-level tokenization, allowing the model to represent both individual symbols and frequently occurring molecular fragments.

The sequence encoder is based on the Transformer architecture [15], which uses self-attention to model long-range dependencies within a token sequence. For molecular SMILES, this mechanism enables the encoder to capture global sequence-level patterns that may correspond to functional groups, branches, rings, and other structural information implicitly represented in the string. Each Transformer layer contains a multi-head self-attention module and a feed-forward network, with residual connections and layer normalization applied to stabilize training.

To obtain the 1D molecular representation, we use ChemBERTa as the pretrained SMILES encoder [2]. ChemBERTa was pretrained on large-scale unlabeled molecular data from ZINC [9] using self-supervised learning, and the model used in this work contains six Transformer layers and twelve attention heads. In TMCA, the parameters of ChemBERTa are frozen during downstream training, so the 1D branch serves as a stable pretrained feature extractor. The output embedding from this branch is denoted as *F*_*s*_, which is later used as the sequential modality representation in the graph-guided cross-attention fusion module.

### 3.2 2D Graph Encoder

The 2D graph encoder (Fig. 1(b)) follows the Graph Attention Network (GAT)-based design used in our previous DMCA framework [27]. Each molecule is represented as a graph, where atoms are nodes and chemical bonds are edges. The encoder consists of four GAT layers [16], with normalization, nonlinear activation, and dropout applied after each layer. After the final GAT layer, we apply both mean pooling and max pooling over the node representations and concatenate the pooled vectors to obtain the graph-level embedding *F*_*g*_.

Different from DMCA, where the graph embedding is used for dual-modality graph–SMILES fusion, TMCA uses *F*_*g*_ as the central query representation in two graph-guided cross-attention modules. This design allows the trainable 2D graph branch to retrieve complementary information from the frozen pretrained 1D sequence encoder and 3D geometric encoder.

### 3.3 3D Geometry Encoder

We further incorporate 3D molecular conformations as the third modality. A molecular conformation describes a molecule through atom types and their corresponding atomic coordinates, providing geometric information such as molecular shape, relative atom positions, and spatial relationships between atoms. These spatial features are not explicitly encoded in either 1D SMILES strings or 2D molecular graphs, but they can be important for molecular property prediction. For this modality, we adopt Uni-Mol as the 3D geometric encoder [28]. Uni-Mol is a pretrained Transformer-based model designed for molecular conformation representation learning. Given atom types and 3D atomic coordinates as inputs, it learns conformation-aware molecular representations by modeling both atom-level features and pairwise spatial relationships. Compared with sequence-based or graph-based encoders, Uni-Mol directly uses coordinate information and therefore provides a complementary geometric view of the molecule.

In TMCA, Uni-Mol is used as a pretrained 3D feature extractor. During downstream training, its parameters are frozen to preserve the geometric knowledge learned during pretraining and to reduce the number of trainable parameters. The output embedding from Uni-Mol is denoted as *F*_*c*_, representing the 3D conformation feature of the molecule. This embedding is then used as the geometric modality input for the tri-modality fusion module described in the following section.

### 3.4 Triple-Modality Cross-Attention Encoder

After obtaining the modality-specific embeddings from the three encoders, TMCA applies two cross-attention modules to learn complementary information among 1D sequential, 2D topological, and 3D geometric representations. More specifically, we use the trainable 2D graph embedding *F*_*g*_ as the central query to integrate information from both the sequential and geometric modalities. Fig. 1(c) shows the first cross-attention encoder between the 2D graph representation and the 1D SMILES representation, where the graph embedding *F*_*g*_ is used to compute the query matrix and the SMILES embedding *F*_*s*_ is used to compute the key and value matrices as follows.

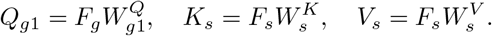

The graph-guided sequence fusion representation is then computed as

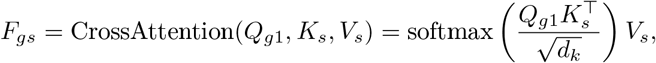

which captures the sequential information from SMILES that is relevant to the graph representation.

The second cross-attention encoder (Fig. 1(d)) handles the 2D graph representation and the 3D geometric representation in a similar fashion. The graph embedding *F*_*g*_ is used to compute the query matrix, and the 3D conformation embedding *F*_*c*_ is used to compute the key and value matrices.

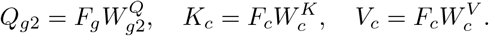

The graph-guided geometry fusion representation is then computed as

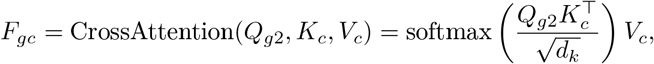

which is the representation learned by using the 2D graph modality to attend to the 3D geometric modality.

After the two cross-attention encoders, the fused representations *F*_*gs*_ and *F*_*gc*_ are combined to form the final tri-modality molecular representation. We consider two fusion strategies: element-wise addition and feature concatenation:

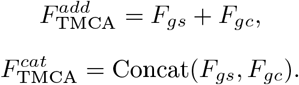

The concatenation strategy preserves information from the graph-guided sequence and graph-guided geometry fusion branches separately, while the addition forces the two fused embeddings into a shared representation space. Both strategies will be evaluated in our experiments. The final prediction is obtained by passing this representation into a multilayer perceptron:

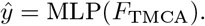

### 3.5 Implementation

We implement TMCA using PyTorch. The 1D sequence encoder is based on ChemBERTa and outputs 767-dimensional SMILES embeddings, which is then used as the default dimensionality for all three modalities. The output from the 3D geometric encoder Uni-Mol is projected to 767 dimensions. The 2D graph encoder is trained from scratch and also produces a 767-dimensional graph embedding. The cross-attention modules use 13 attention heads. Cross-entropy loss and the Adam optimizer are used for model training. For the final fusion step, we evaluate both concatenation and element-wise addition strategies. Furthermore, for the ablation study, we implement model variants that unfreeze the pretrained 1D sequence encoder and 3D geometric encoder. So in total, four model variants are implemented and they are denoted as TMCA-FC, TMCA-FA, TMCA-UC, and TMCA-UA, for frozen+concatenation, frozen+addition, unfrozen+concatenation, and unfrozen+addition, respectively.

## 4 Experiments

To evaluate the effectiveness of the proposed TMCA framework, we conduct molecular property prediction experiments on classification tasks from the MoleculeNet benchmark datasets [22]. We briefly introduce the state of the art competing approaches, the datasets, and the experimental design in this section.

### 4.1 Baseline approaches

We compare TMCA with representative molecular property prediction methods based on single-modality as well as those based on multi-modalities. The single-modality baselines include graph-based methods, such as Hu et al. [8] and MolCLR [18], and a SMILES-based method, ST [7]. We also include geometry-enhanced graph pretraining methods, including GEM [3] and GraphMVP [13], both of which incorporate 3D geometric information during representation learning or pretraining. For models based on two modalities (SMILES and graph representations), we include DVMP [29], GraSeq [5], and DMCA [27]. For multi modality models, we include MSSGAT [26], which integrates graph, substructure, and fingerprint-related molecular features, and SGGRL [19], which combines sequence, graph, and geometry modalities. Most baseline results are obtained from their corresponding publications. SGGRL is rerun under our experimental setting because it is the only other approach that also utilizes all three modalities. The hyperparameter settings used for rerunning SGGRL are summarized in the Appendix (Table A1).

### 4.2 Datasets

We evaluate TMCA and compare its performance with other competing approaches on four datasets from MoleculeNet [22]: BBBP, BACE, ClinTox, and HIV. A brief description of these datasets can be found in our earlier paper [27]. BBBP, BACE, and HIV contain binary labels. Although ClinTox originally contains two labels, our previous analysis in DMCA [27] has shown that these two labels were highly correlated. Therefore, we only use one label for ClinTox and all the prediction are binary classification tasks. For baseline methods that report two-label ClinTox results, we use the average of their performance over the two labels for comparison.

### 4.3 Experiment Design

For each molecule, the 1D SMILES representation is directly obtained from MoleculeNet. The 2D molecular graph is generated from the SMILES string using RDKit [10]. We generate 3D conformations using the Uni-Mol conformation generation pipeline. Molecules for which valid 3D conformations could not be generated were excluded from the experiments. We then applied scaffold splitting to the remaining molecules, using a training/validation/test ratio of 0.8/0.1/0.1. Scaffold splitting is generally more challenging than random splitting because it evaluates the model on molecular scaffolds that are different from those seen during training [8, 27]. Each experiment was repeated three times with different random seeds. We use the Area Under the Receiver Operating Characteristic Curve (AUC-ROC) as the main evaluation metric. The mean and standard deviation of AUC-ROC were recorded.

Furthermore, we consider four different variants of TMCA, as mentioned earlier, which examine the effects of different strategies for the final fusion step and different training strategies. All experiments are conducted on the high-performance computing platform at the Ohio Supercomputer Center.

## 5 Results

### 5.1 Comparison with baselines

Table 1 reports the AUC-ROC results on four MoleculeNet classification datasets of all approaches. TMCA-UC achieves the best AUC-ROC on BBBP and TMCA-FC achieves the best result on ClinTox. DVMP obtains the best result on BACE, and MolCLR achieves the best result on HIV. This suggests that molecular property prediction remains dataset-dependent, and no single architecture dominates across all tasks. Because no single method consistently achieves the best AUC-ROC across all datasets, we follow the rank-score evaluation strategy used in DMCA [27] to summarize the overall performance, which is shown in Fig. 2. In this figure, we use results from TMCA-FC to represent the performance of TMCA.

**Table 1.**
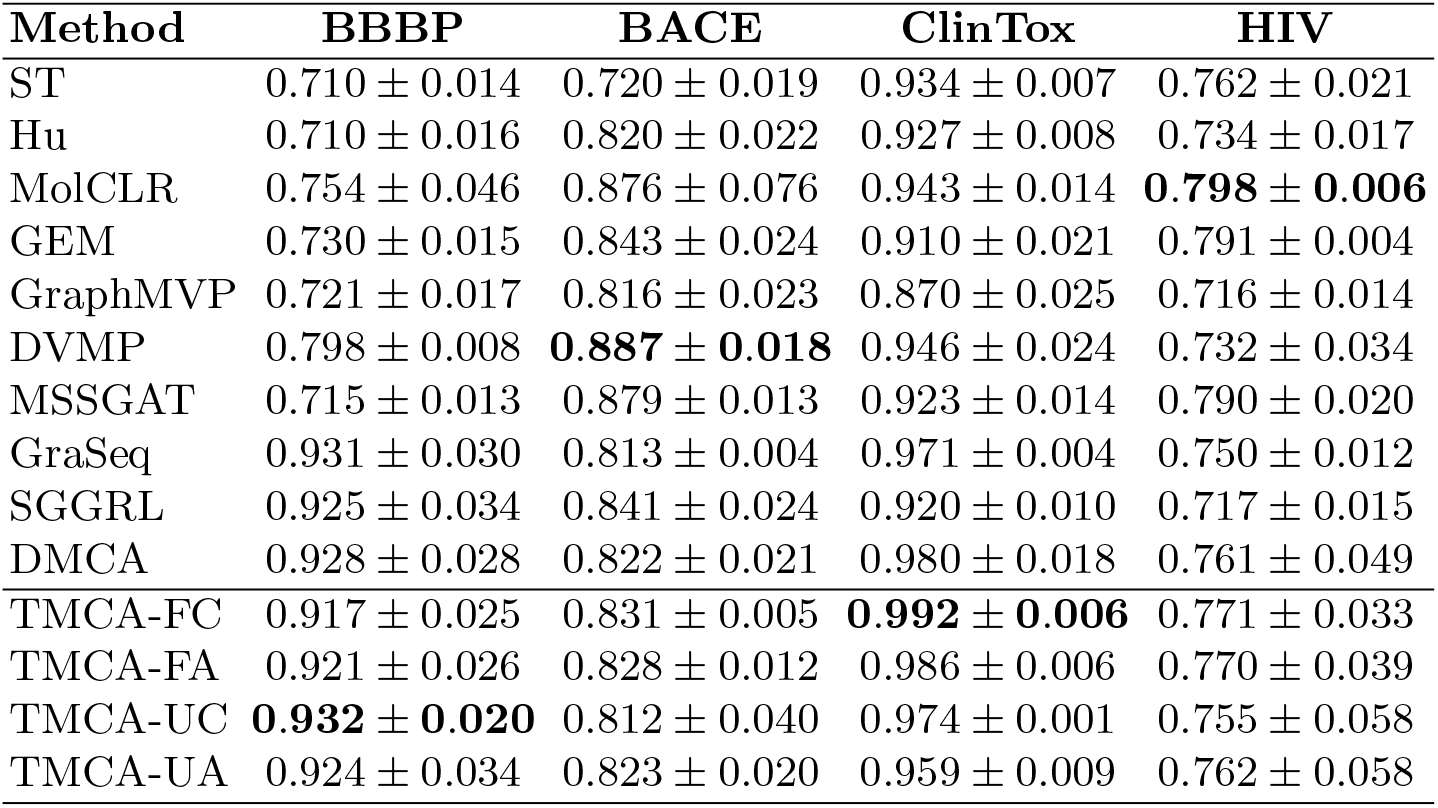
AUC-ROC performance comparison on four MoleculeNet classification datasets. The best result for each dataset is shown in bold.

**Fig. 2.**
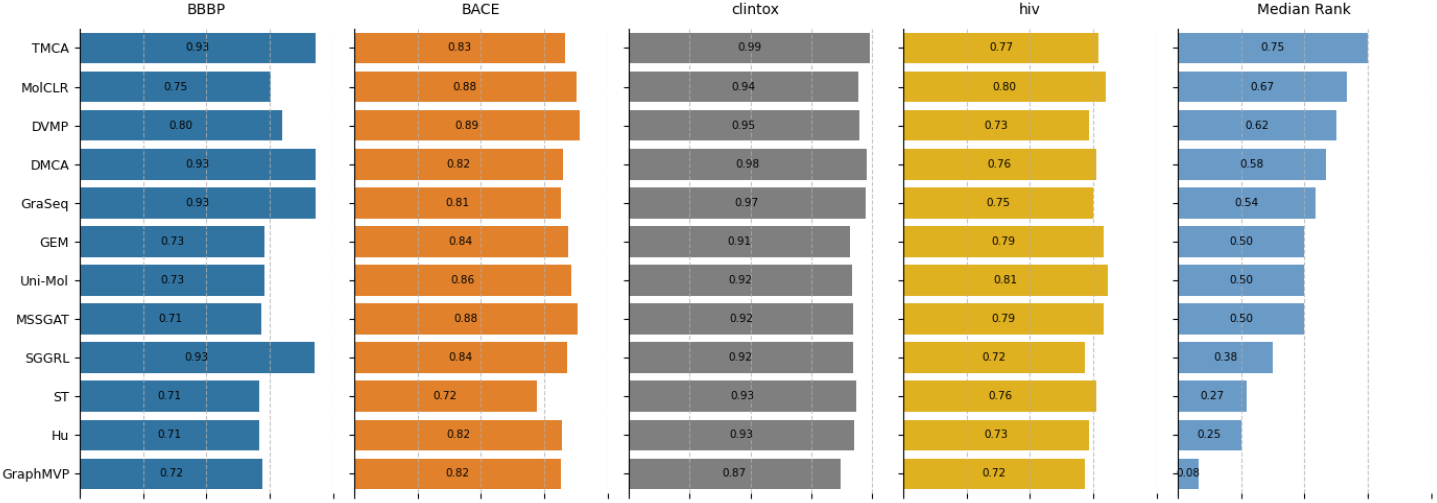
Overall ranking scores of all compared methods across the MoleculeNet classification datasets. Higher scores indicate better overall performance after dataset-wise min-max normalization.

The ranking in Fig. 2 shows that TMCA achieves the strongest overall performance among all the compared methods, which demonstrates the effectiveness of the proposed tri-modality learning framework. By using the 2D graph branch as the central query modality and retrieving complementary information from pretrained 1D sequence and 3D geometric encoders, TMCA effectively integrates sequential, topological, and geometric molecular information for molecular property prediction.

### 5.2 Ablation Study

Based on the results from the four variant models Table 1, the frozen-encoder variants generally provide more stable performance across datasets with lower standard deviations in most cases comparing with the unfrozen counterparts. In particular TMCA-FC achieves the best result on three datasets (ClinTox, BACE, and HIV) among the four variants. TMCA-FA shows similar behavior with slightly lower performance than TMCA-FC on BACE, ClinTox, and HIV. These results suggest that freezing the pretrained ChemBERTa and Uni-Mol encoders can preserve useful sequence and geometric knowledge learned during pretraining, while allowing the trainable 2D graph encoder and cross-attention modules to adapt to downstream tasks.

In terms of the fusion strategies, concatenation performs better than addition in the frozen setting. TMCA-FC outperforms TMCA-FA on BACE, ClinTox, and HIV, while TMCA-FA is only slightly better on BBBP. This indicates that keeping the graph-guided sequence representation and graph-guided geometry representation as separate feature components is generally more effective than directly merging them through element-wise addition. In the unfrozen setting, TMCA-UC achieves better results on BBBP and ClinTox comparing with TMCA-UA. It has lower performance on BACE and HIV but with large variance. The behavior is more likely due to limited training time.

Overall, the ablation results support the main design of TMCA. Using frozen pretrained 1D and 3D encoders together with a trainable 2D graph encoder provides a stable way to combine pretrained and non-pretrained representations. In addition, concatenation is generally more effective than additive fusion in most cases.

## 6 Conclusion and Discussion

In this paper, we propose Tri-Modality Cross-Attention (TMCA), a molecular representation learning framework that integrates 1D SMILES strings, 2D molecular graphs, and 3D molecular conformations for molecular property prediction. TMCA uses pretrained encoders for the 1D sequence and 3D geometric modalities, and trains a 2D graph encoder from scratch as the central query branch in the cross-attention fusion module. By learning graph-guided sequence and graph-guided geometry representations, TMCA provides an effective way to combine pretrained and non-pretrained molecular encoders. We evaluated TMCA on four MoleculeNet classification datasets and compared it with representative single-modality and multimodal baselines. The results show that TMCA achieves strong overall performance. The ablation study suggests that freezing the pretrained encoders and using concatenation fusion generally provides better and more stable results with fast training. One limitation of TMCA is that it requires valid 3D conformations, so molecules with failed conformation generation are excluded from the experiments. In addition, because most baseline results are taken from published papers, differences in preprocessing and data splitting may affect direct comparability. Limited number of repeated experiments due to constraints from computational resources makes it hard to assess the significance of performance difference. Future work will extend TMCA to handle missing-modality cases where reliable 3D conformations may not always be available and perform more experiment studies on more datasets and more prediction tasks.

## Acknowledgments

This work is supported by the National Science Foundation (grant Nos. CCF-2200255, CCF-2006780, IIS-2027667) and the National Institutes of Health (grant Nos. U01AG073323, R01HG009658). This work was supported in part by an allocation of computing time from the Ohio Supercomputer Center.

## Appendix

**Table A1.**
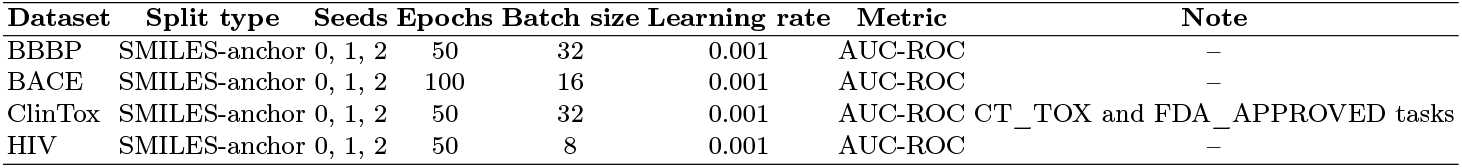
Hyperparameter settings used for rerunning SGGRL.

